# The sensitivity of acute myeloid leukemia to CDK8/19 inhibitors is determined by their metabolic profile

**DOI:** 10.64898/2025.12.14.694205

**Authors:** Ekaterina A. Varlamova, Daria V. Andreeva, Alexandra A. Dalina, Alexandr V. Ivanov, Artemy P. Fedulov, Elena V. Misnik, Natalia G. Pavlenko, Alexandra V. Bruter, Victor V. Tatarskiy

## Abstract

**Background:** Cyclin-dependent kinases CDK8/19 are serine/threonine kinases that regulate transcription as part of the Mediator complex and phosphorylate several non-transcriptional substrates in both normal and tumor cells. Several studies have demonstrated that inhibition of CDK8/19 leads to selective cell death in acute myeloid leukemia (AML) cells with favorable adverse effect profile. However, the exact mechanism of AML sensitivity to CDK8/19 inhibitors (CDK8/19i) is poorly understood. One of the key goals of the current research was to identify the molecular mechanisms underlying this sensitivity.

**Methods:** AML cell lines were stratified by CDK8/19i and CCNC KO sensitivity using the DepMap database and published cytotoxicity data. Mean KO dependency scores grouped by Hallmark categories were correlated with CDK8/19i sensitivity (Spearman), and metabolic indices were calculated as mean log2(TPM+1) expression of pathway genes. Viability of AML lines MV4;11, KG-1, THP-1, Kasumi-1 was assessed after 120 h CDK8/19i (SenB and SNX631) via resazurin or flow cytometry cell cycle analysis. RNA sequencing of all AML lines after 72 h with 1 μM SenB was analyzed with DESeq2 (p_adj < 0.05). Oxygen consumption rate of all AML lines was measured after 72 h with 1 μM SNX631 using Seahorse XF Mito Stress Test or Seahorse XF Glyolysis Stress Test (Two-way ANOVA, p < 0.05). Intracellular metabolite profiling of all four AML lines after 72 h treatment with 1 µM SNX631 was performed by GC-MS, with peak areas normalized to total protein content. Metabolites with p_val < 0.05 and |log FC| ≥ 0.6 (Welch’s two-sample *t*-test) were considered significant.

**Results:** According to the DepMap database CDK8/19i sensitivity is associated with sensitivity to knockout of metabolic-associated genes. Resazurin assay ranked sensitivity as KG-1, MV4;11 > THP-1 > Kasumi-1 for both inhibitors. Cell cycle analysis after 120 h showed that inhibition of CDK8/19 caused induction of cell death in MV4;11, while KG-1 cells exhibited reduced metabolic activity. GlycoStress assay showed that the MV4;11 cells which are the most sensitive to CDK8/19i have the highest glycolytic capacity. RNA-seq analysis showed that CDK8/19i after 72 h downregulated glycolysis genes only in sensitive KG-1 and MV4;11. MitoStress parameters are reduced most in MV4;11, then KG-1/THP-1, but not in resistant Kasumi-1. CDK8/19i led to decrease in glutaminolysis metabolites only in sensitive KG-1 and MV4;11 cells.

**Conclusion:** CDK8/19i in AML predominantly targets cells with a glycolytic-active metabolic phenotype, leading to changes in metabolic composition, disruption of TCA, reduced expression of glycolysis-related genes and decreased glycolysis. Our data suggest that the metabolic profile of AML cells may serve as a functional marker for identifying tumors most likely to respond to CDK8/19i.

## Introduction

Acute myeloid leukemia arises from malignant clonal expansion of myeloid progenitors in their differentiation block[1]. AML is an aggressive form of leukemia that primarily affects older adults; the median age at diagnosis is approximately 70 years[2]. Although the survival rate of patients under 60 years of age has increased from 13% to 55% over the past 50 years, in older patients with comorbidities this parameter has only increased from 8% to 17%[3].

CDK8/19 inhibitors represent a promising therapeutic approach that offers opportunities to overcome the limitations of conventional chemotherapy. CDK8/19 inhibitors are specifically active in AML and prostate cancer and demonstrated high efficacy against relapsed and refractory AML and good tolerability[4–6]. The most clinically advanced CDK8/19 inhibitor, RVU120, is currently investigated in Phases 1b and 2 for AML. However, it must be noted that not all AML tumors are sensitive to CDK8/19i and there are no accepted markers to discriminate sensitive ones. A combination of RVU120 with venetoclax (an inhibitor of an anti-apoptotic protein Bcl-2) for patients with AML is also being studied; however, despite the synergism, multiple side effects were noted[7–9]. Additional pre-clinical research showed that CDK8/19i combination with BCR-ABL inhibitors such as imatinib has been proposed to overcome chemoresistance in chronic myeloid leukemia[10].

A study by Pelish et al. showed that pharmacological inhibition of CDK8/19 leads to AML cell death due to transcriptional dysregulation, primarily of genes regulated by super-enhancers, but a specific mechanism was not identified[4]. Although current studies indicate a role for phosphorylation STAT1 and STAT5 at serine 727 as a marker of AML cell sensitivity, they are not universal and cannot act as clinical markers [11,12]. Recent research showed that CDK8/19 function has been demonstrated in lipogenesis [13], cellular response to hypoxia [14], glycolysis [15] and steroid biosynthesis [16], confirming the important role of these kinases in cellular metabolism. In this study, we observed that the sensitivity of AML cells to CDK8/19i correlated with their basal glycolytic activity.

Here we propose that sensitivity to CDK8/19i is determined by a specific metabolic alteration in AML cells, caused by CDK8/19 inhibition. It has been shown that CDK8/19 play important roles in metabolism and cell senescence. In this study, we analyzed open-source DepMap databases and examined the transcriptome, metabolome, OCR and ECAR of AML cells with varying sensitivities to CDK8/19i before and after exposure to CDK8/19i. We found a correlation between AML cell sensitivity to CDK8/19i and sensitivity to KO of genes associated with mitochondrial function, glycolysis, and lipid metabolism, and changes in these processes which occur only in sensitive cells.

## Material and methods

### Compounds

CDK8/19 inhibitors (CDK8/19i) Senexin B (SenB) was from Biocad, Russia and SNX631 was a gift from Senex Biotechnology (Columbia, SC, USA). Stock solutions (10 mM) were prepared in dimethyl sulfoxide (DMSO) and stored at −20°C. Working dilutions in complete culture medium were prepared immediately before each experiment.

### Cell culture

Human AML cell lines MV4;11, KG-1 (Russian Collection of Cell Cultures, Saint-Petersburg, Russia), THP-1 (TIB-202, ATCC), Kasumi-1 (the cells were provided by the Department of Genetics, Faculty of Biology, Belarusian State University) were propagated in RPMI-1640 (PanEco, Moscow, Russia) with 10% fetal bovine serum (Biosera, Cholet, France), 2 mM L-glutamine, 100 U/mL penicillin and 100 μg/mL streptomycin (PanEco) at 37°C and 5% CO2 in humidified atmosphere.

### Cell viability assay (Resazurin assay)

Cells (75×10^3^ cells/mL) in the logarithmic phase of growth were plated into 96-well plates (SPL, Korea) and treated with CDK8/19i for 120 h at indicated concentrations. Wells containing only DMSO served as controls. Control wells contained 0.02% DMSO. To assess cells proliferation, resazurin (Sigma Life Sciences, Canada) was added to the wells at a final concentration of 0.5 mM and incubated for another 3 h, after which the fluorescence (Ex/Em = 560/620 nm) in each well was measured using Tecan Infinite M200 (Tecan Group Ltd, Männedorf, Switzerland).

### Cell cycle assay

Cells were plated as described above in 10 cm Petri dishes (SPL, Korea) and incubated in the presence of inhibitors for 120h. Cells with an equivalent amount of DMSO served as a control. At the end of the incubation, cells were fixed with 100% ice-cold methanol for 10 minutes at -20°C, washed in PBS (Eco-service, Russia), and stained with DAPI solution (5 μg/ml) (Thermo Fisher Scientific, Waltham, MA). Cell cycle phase distribution was assessed by flow cytometry analysis on a CytoFlex 26 (Beckman Coulter, Indianapolis, IN) in PB-450 channel.

### Immunoblotting

Protein lysis and western blotting were performed as described in[6]. Cells were lysed in RIPA buffer supplemented with protein inhibitor cocktail (Sigma-Aldrich, St. Louis, MO). Total protein concentration was quantified by the Bradford method. Absorbance at 570 nm was measured with a Tecan Infinite 500 Plate Reader (Tecan Trading AG, Switzerland). Proteins were separated by SDS-PAGE and transferred onto 0.45 μm nitrocellulose membrane (Bio-Rad, Hercules, CA). After blocking with 5% skimmed milk, membranes were treated with primary antibodies against CDK8 (Cat #17395, Cell Signaling Technology), CDK19 (described in [17]), CCNC (Cat #68179, Cell Signaling Technology), β -actin (Cat#A2228, Sigma) and incubated at 4°C overnight. Membranes were washed with Tris-borate saline with Tween 20 (TBS-T) and incubated for 1 h at room temperature with secondary anti-rabbit IgG (Cat#7074, Cell Signaling Technology) or anti-mouse IgG (Cat #7076, Cell Signaling Technology) antibodies. Membranes were visualized with the Clarity Western ECL Substrate (Bio-Rad) using an iBright FL1500 Imaging System (Invitrogen, Waltham, MA).

### Seahorse Extracellular Flux Assay

To investigate the effect of CDK8/19i on the energy balance of AML cells, we performed the extracellular flux assay after 72h treatment with 1 μM SNX631. This time point was chosen as an intermediate point when significant cell death was not yet observed based on the dose-dependent cell survival data. Cells, incubated for 72h in the presence of equimolar concentrations of DMSO, served as a control group.

To perform the analysis, MV4;11 (500,000 cells per well), Kasumi-1 (800,000 cells per well), THP-1 (600,000 cells per well), and KG-1 (700,000 cells per well) cells were seeded into the Seahorse XF24 V7 PS Cell Culture Microplates (Agilent Technologies, USA) coated with poly-L-lysine. Cells were centrifuged at 200 g for 5 min and incubated in 500 μl of the Seahorse Assay Medium supplemented with 2 mM glutamine and 10 mM glucose for the MitoStress test for 30 minutes at 37°C in the absence of CO_2_. The measurements were performed following the standard protocol (3 min mix, 2 min wait, 3 min measure) at 37°C after plate equilibration, with 3 measurement cycles between injections. For the GlycoStress test, cells were injected with 10 mM glucose (AppliChem), 2 μM oligomycin (Sigma Aldrich), and 50 mM 2-deoxyglucose (2DG). For the MitoStress test, cells were injected with 2 mM oligomycin (Sigma Aldrich), 1 μM FCCP (Sigma Aldrich), and the combination of 1 μM rotenone and 2 μM of antimycin A (Sigma Aldrich). Results were presented as OCR (oxygene consumption rate) per 1000 cells.

### RNA sequencing and analysis

AML cells (3×10^5^ cells/mL) were treated with the vehicle (0.02% DMSO) or 1 μM SenB for 72 h. Total RNA was extracted with RNeasy Mini Kit (QIAGEN, Netherlands). RNA sequencing libraries were prepared using a TruSeq Stranded mRNA kit (RS-122-2101, Illumina, United States) according to the manufacturer’s protocol. The cDNA amount was determined using a 4150 TapeStation system (Agilent, United States). Sequencing was carried out using an Illumina NextSeq 2000 sequencing system (100 cycles). Quality of raw sequencing data was checked using the FastQC tool (v. 0.11.9). All reads were aligned to the GRCh37 reference human genome with the use of the STAR aligner (v. 2.604a) and counted using featureCounts (v. 1.6.4). Data convergence was assessed using principal component analysis (PCA) (Supplementary Fig. S1). Differentially expressed genes (DEGs, log2FC > 1.5 and p adjusted for multiple comparisons (p adjust) < 0.05) were identified using the DESeq2 package in the R language. Gene Ontology (GO) enrichment analyses were carried out using the clusterProfiler package in the R language. The ggplot2, pheatmap and clusterProfiler R libraries were used for data visualization.

TF activity was inferred using the decoupleR R package and the human COLLECTRI network. For each AML cell line, DESeq2 gene-level statistics were used as input for the ULM method. For each TF, we computed activity scores, BH-adjusted P values, and the number of significantly and consistently regulated target genes (padj < 0.05, |log2FC| ≥ 0.5, regulation direction concordant with interaction sign). Network analysis was performed with the STRING database [18].

### Metabolite Quantification by Gas Chromatography-Mass Spectrometry

Metabolite quantification was performed by the GC-MS method described in[19]. Cells were plated as described above in 10 cm Petri dishes (SPL, Korea) and incubated in the presence of SNX631 for 72 h. Cells with an equivalent amount of DMSO served as a control. After incubation cells were centrifuged at 200 g for 5 min and the pellets were freezed at −80 °C prior to further processing. Total protein was quantified with the Micro BCA kit (Thermo Fischer Scientific, Waltham, MA, USA). Measurements were carried out on a gas chromatography-mass spectrometer coupled with a monoquadrupole mass spectrometric detector (Crystal 5000, Yoshkar-Ola, Russia). Samples (1 μL) were injected into a helium stream at a rate of 1 mL/min; electron ionization was performed at 70 eV. Metabolites were separated on an HP-5 ms column (30 m × 0.25 mm × 0.25 μm) (Agilent Technologies, Santa Clara, CA, USA). The injector, transfer line, and ion source temperatures were set to 250 °C, 290 °C, and 230 °C, respectively. Raw spectra were analyzed by Chromatec Analytic Software v.3.0.0.2 (Yoshkar-Ola, Russia). Quantification of the compounds was performed in the single ion monitoring mode in each channel, determining for each metabolite a characteristic ion in a narrow range of retention times for a given compound. Calculated peak areas were filtered by pooled QC samples with 30% RSD cutoff with following normalization to the signal of Norvaline as an internal standard. Additionally, peak areas were normalized to the protein concentrations in samples.

Principal component analysis (PCA) was performed on the autoscaled data matrix using the prcomp function in R to assess sample clustering and identify potential outliers (Supplementary Fig. S2). Replicate outliers were identified visually from PCA score plots and excluded prior to differential analysis, retaining three to four biological replicates per group. Differential abundance between SNX631-treated and vehicle control groups was assessed separately for each cell line using two-sided Welch’s t-tests. Metabolites with |log FC| ≥ 0.6 and p < 0.05 were considered significantly altered.

### Open-source data

To separate cells based on sensitivity to CDK8/19i, we collected IC50 values for known CDK8/19 inhibitors (CDK8/19i) (CA, RUV120, SenexinB, MK256) for various AML cell lines from open-source articles and patents (Supplementary Table 1). The minimal IC50 for each cell line (minIC50) was used to classify cells as very sensitive (minIC_50_ < 100 nM), sensitive (minIC_50_ < 700 nM) and non-sensitive cells (minIC_50_ > 700 nM) to CDK8/19i. The OCI-AML3 cell line was isolated as a separate group due to its intermediate IC50 values (∼200 nM). For CRISPR_CCNC and RNAi_CCNC, cells were also divided into groups based on sensitivity and resistance to knockout and siRNA, respectively. For CRISPR_CCNC, groups were defined as very sensitive (< -0.8), sensitive (< -0.5), and non-sensitive (other). For RNAi_CCNC, groups were defined as very sensitive (< -0.6), sensitive (< -0.3), and non-sensitive (other).

We obtained data on the impact of gene knockout (KO) on cell survival from the DepMap database (CRISPR Chronos) and grouped them using MSigDB (Hallmark database) for the corresponding biological processes and calculated the average values for each group using the matrixStats package in R. For all groups, the values were normalized (z-scale). Data from the DepMap CRISPR (DepMap Public 25Q3+Score, Chronos) were analyzed using the matrixStats library in R, and the correlation with the CDK8/19i minIC50 was assessed using the Spearman method. To evaluate the metabolic profile of AML cells depending on their sensitivity to CDK8/19i using gene expression data associated with various metabolic pathways we used a database of gene expression in various AML cells (DepMap Expression Public 25Q3). Genes were grouped according to their correspondence to Hallmark databases and then calculated the average expression values within each group for each individual cell line.

### Statistical analysis

Statistical analysis performed with GraphPad Prism 10. Statistical significance was defined as p□<□0.05, and the results were annotated as follows: *p□<□0.05, **p□<□0.01, ***p□<□0.001, ****p□<□0.0001; ns: not significant.

## Results

### Sensitivity to CDK8/19 inhibition in AML cells correlates with their sensitivity to knockout of metabolism-associated genes

To address the mechanisms of AML sensitivity to CDK8/19i we first divided AML cell lines into CDK8/19i-sensitive and resistant based on their minimal published IC50 (minIC50) and their sensitivity to *CCNC* gene knockout (CRISPR_CCNC) from the DepMap database (Fig. 1A). Cyclin C was chosen because as a CDK8/19 partner it is strictly required for CDK8/19 activity[20]. CRISPR_CCNC and minIC50 are well aligned (Fig. 1A). We performed further analysis for AML cell lines for which both CRISPR_CCNC and IC50 values were known.

**Figure 1.**
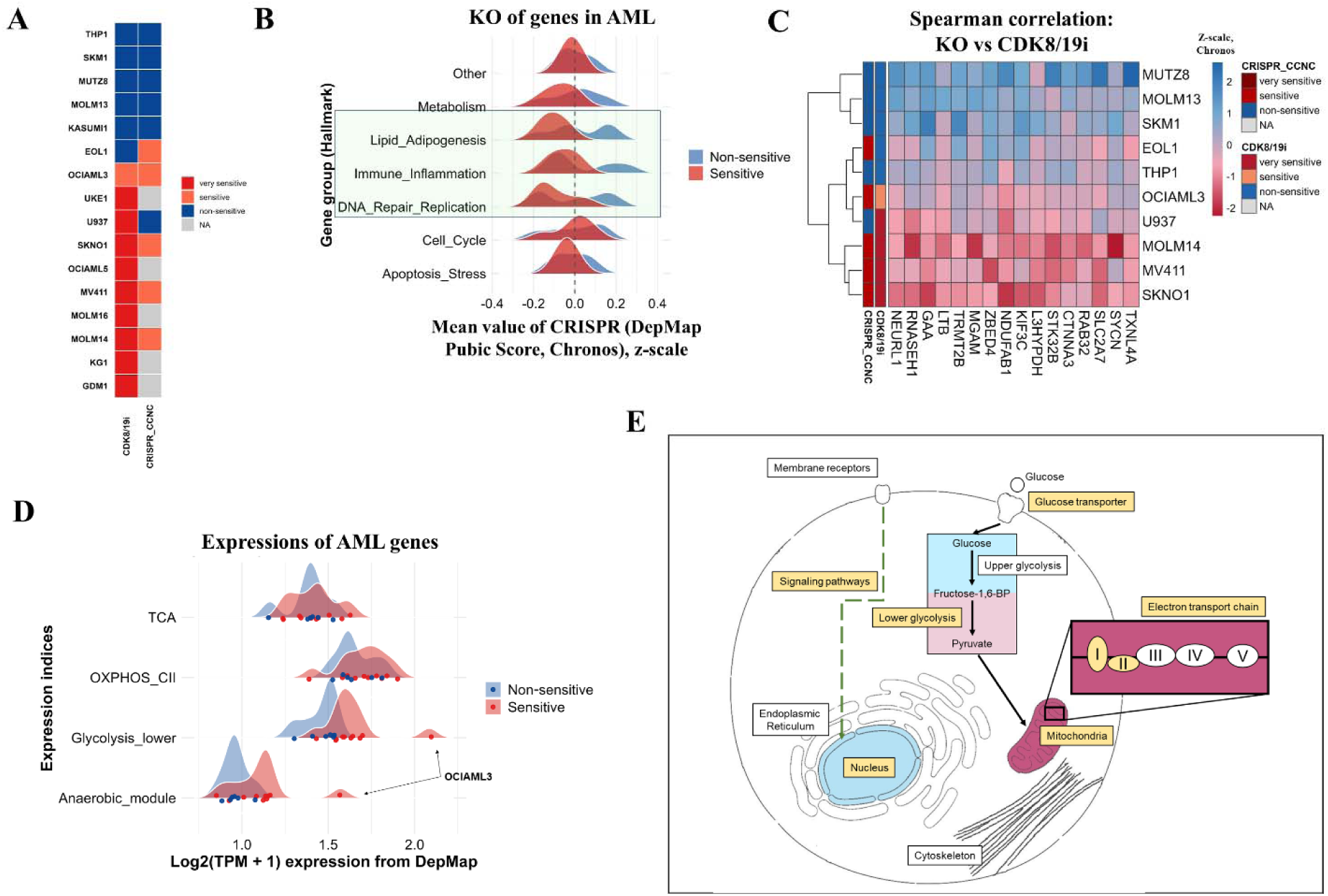
AML cell sensitivity to CDK8/19i correlates with their metabolic properties. **A.** AML cell lines and their sensitivity to CCNC KO (CRISPR_CCNC (DepMap Public 25Q3+Score, Chronos)) and as determined by DepMap or CDK8/19i as determined by minIC50 from published data. **B.** Distribution ridgeplots of CDK8/19i sensitive and resistant AML cells based on their average sensitivity to gene KO (DepMap, Chronos) in the groups determined using the Hallmark database. Data transformed using z-scaling. Groups of KO with the greatest differences between groups of cells are marked in green. **C.** Heatmap of genes whose KO directly correlates with AML sensitivity to CDK8/19i. Data was obtained from the DepMap (Public 25Q3+Score, Chronos) using Spearman correlation. Data were transformed using z-scaling. **D.** Distribution ridgeplots of CDK8/19i sensitive and resistant AML cells based on their metabolic indices (genes obtained from DepMap Expression Public 25Q3 database). The indices with the greatest differences between CDK8/19i sensitive and resistant AML cells are presented. **E.** A schematic diagram summarizing the major cellular processes that may be involved in the sensitivity of AML cells to CDK8/19i.

Our first goal was to identify genes such that sensitivity to their knockout (KO), either individual or combined, could divide cell lines in the same way they are divided by sensitivity to CDK8/19i and CRISPR_CCNC. That could mean that the products of these genes are involved in those CDK8/19 dependent processes which confer AML cell lines their sensitivity. First, we identified three functional groups of genes with correlation between mean KO score for the genes in the group and AML sensitivity to CDK8/19i: lipid metabolism and adipogenesis, immune response and inflammation, DNA replication and repair (Fig. 1B). Next, we identified specific genes from the processes mentioned above that contribute most to the correlation (Supplementary Fig. S3A): those were genes related to mitochondria functioning, including mitochondrial RNA translation and mitochondrial ribosome synthesis, the Krebs cycle and the ETC, fatty acid oxidation, steroid metabolism and lipogenesis. Interestingly, genes encoding some subunits of RNA polymerases I, II, and III (*POLR1C* and *POLR2F*) and genes encoding proteins required for transcription elongation and pausing (*NELFB*, *SUPT5H*) also contributed to the pathways mentioned above. These data indicate that the sensitivity of AML cells does not depend on any specific target, but on a group of genes regulating similar processes.

Alternatively, we searched for genes whose individual KO sensitivity directly correlated with cell sensitivity to CDK8/19i (Fig. 1C). We identified 16 genes including those associated with the regulation of mitochondrial gene translation (*TRMT2B, RNASEH1*), nuclear gene transcription (*TXNL4A*, *ZBED4*), oxidative phosphorylation (OXPHOS) (*NDUFAB1*), glucose and amino acid metabolism (*SLC2A7*, *SYCN*, *L3HYPDH*, *GAA*, *MGAM*). Besides these genes we also identified genes from Wnt/b-catenin (*CTNNA3*), TNFa (*LTB*) and NOTCH (*NEURL1*) signaling pathways and one gene associated with cell adhesion (*KIF3C*).

As we spotted the correlation between sensitivity to the metabolic genes KO and CDK8/19i we further used the DepMap expression data to calculate transcriptomic metabolic indices for distinct parts of cellular metabolism. We identified four classes of genes with notable differences: the Krebs cycle (TCA), the second complex of the ETC (OXPHOS_CII), the second part of the glycolytic process from fructose-1,6-biphosphate to pyruvate including ATP synthesis (Glycolysis_lower), and anaerobic respiration (the combination of glycolysis, lactate synthesis and PDK coding genes) (Anaerobic_module) (Fig. 1D, Supplementary Fig. S3B). We next performed clustering of AML cell lines based on the values of these 4 gene groups and found that the average expression of selected genes had a correlation with both *CCNC* KO and CDK8/19i values, which indicates a common metabolic pattern of CDK8/19i-sensitive cells (Supplementary Fig. S3C).

Thus, we determined that for the sensitivity of cells to the CDK8/19i, cellular processes associated with glycolysis, the first and second complex of OXPHOS, and processes associated with transcription play an important role (Fig. 1E, yellow highlights).

### CDK8/19 are required for metabolic genes expression of AML cells

To confirm our hypothesis about the role of metabolism in the sensitivity of AML cells to CDK8/19i, we next performed several experiments with available AML cell lines. In this study, two approaches were used to classify AML cells as sensitive or resistant following 120 h incubation with CDK8/19i (Senexin B (SenB) and SNX631) – cell viability assessment using the resazurin staining and analysis of cell cycle phase distribution. We first confirmed that MV4;11 and KG-1 cells are sensitive, while THP-1 and Kasumi-1 cells are resistant to CDK8/19i (Fig. 2A). In the resazurin test KG-1 cells appeared to be the most sensitive (IC_50_ is 0.15±0.07 µM for SNX631and 0.37±0.1 µM for SenB), MV4;11 has moderate sensitivity (IC_50_ is 0.31±0.04 µM for SNX631 and 1.2±0.2 µM for SenB), while Kasumi-1 were the most resistant. At the same time cell cycle analysis CDK8/19 inhibition in KG-1 cells lead only to modest increase in cell death (up to 20%) after 120 h, while MV4;11 cells exhibited greater cell death rate (subG1 up to 60%) (Fig. 2B). It is possible that the reduction of resazurin to resorufin is affected by CDK8/19i in AML, as it slows down metabolic activity especially in KG-1 cells. There were no differences in the distribution of cells across the cell cycle in resistant Kasumi-1 and THP-1 cells after 120 h of incubation in the presence of both CDK8/19i, and subG1 does not exceed 10%. Interestingly, CDK8/19i induced a low cytostatic effect in THP-1 cells (up to 10% delay in S-phase entry) without apparent cell death, suggesting that the lower IC_50_ observed by resazurin staining reflects reduced proliferation rather than cytotoxicity in this cell line. The discrepancies between the results of assessing the viability of AML cells in the presence of CDK8/19i using a metabolic assay and cell cycle analysis points to the important role of kinases in maintaining active cell metabolism in sensitive AML cells. The sensitivity of AML cells is not determined by either the presence or the level of CDK8 and/or CDK19 kinases (Fig. 2C).

**Figure 2.**
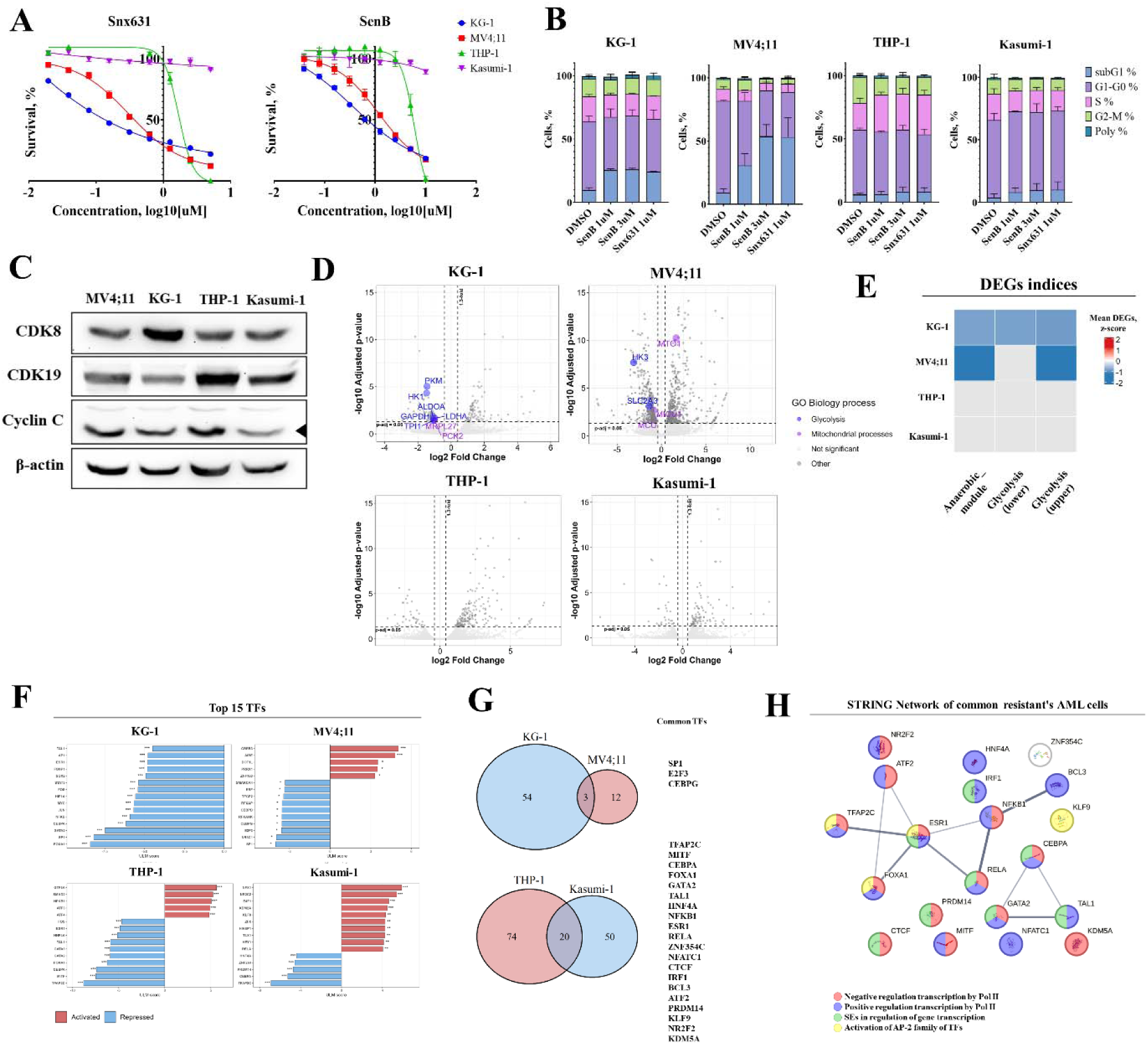
Decreased expression of glycolytic genes occurs only in sensitivity to CDK8/19i AML cells. **A.** Results of assessing the viability of AML cells in the presence of CDK8/19i SenB or SNX631 using resazurin staining after 120h of incubation. Data are presented as mean ± SD, *n* = 3. Both inhibitors are active at submicromolar concentrations against MV4;11 and KG-1 cells, but not THP-1 and Kasumi-1. **B.** Cell cycle distribution after 120 h of treatment with SenB and SNX631. Data are presented as mean ± SD, *n* = 2. С. Western blot of untreated AML cells. Black arrows point to the specific band. **D.** Distribution graphs (Volcano plots) of differentially expressed genes (DEGs) after 72 h of incubation of AML cells in the presence of 1 μM SenB. **E.** Metabolic indices of DEGs that change only in CDK8/19i-sensitive AML cells. Data were transformed using z-scaling. **F.** Top 15 transcription factors (TFs) derived from DEG enrichment by RNA-seq after 72 h in the presence of SenB performed using decoupleR (ULM-score). * – *p*<0.5, ** – *p*<0.1, *** – *p*<0.01, **** – *p*<0.001. **G.** Vienn diagrams of intersecting TFs after enrichment with DEGs. **H.** The STRING network of common TF-enriched resistant AML cells.

To study the transcriptomic changes in the AML cells in response to CDK8/19i, we performed RNA sequencing analysis after 72h in the presence of 1 μM SenB (Supplementary Fig. S4A). Large transcriptomic changes were detected only in MV4;11 (645 DEGs log2 FC > |0.4|, p_adj < 0.05), while in the other cell lines the number of DEGs did not exceed 200 (Supplementary Table 2). GO analysis revealed that SenB resulted in differential expression of genes related to the immune response in all cell lines (Supplementary Fig. S4B). The most altered genes were those related to leukocyte immune response activation and differentiation in all cell types. Among all DEGs, we did not find any genes that changed consistently and similarly in all four lines (Supplementary Fig. S5), however, the molecular processes affected were similar.

As expected from open-source data analysis (Fig. 1) only sensitive AML cell lines showed changes in the expression of genes related to metabolism (Fig. 2D). In MV4;11 and KG-1 cells we observed a decrease in the expression of the hexokinase genes *HK3* and *HK1* (log2FC = -3.121 and -1.474, respectively). Hexokinases phosphorylate glucose at the first stage of glycolysis. Expression of genes encoding enzymes of acetate metabolism and oxidative stress *ACSS2* (log2FC = -0.831), *ALDH3A2* (log2FC = -0.75), *ALDH3B1* (log2FC = -1.122) was also reduced in both sensitive cell lines. In KG-1 cells, we additionally detected a decrease in the expression of the pyruvate kinase PKM gene (log2FC = -1.447), which performs the last step of glycolysis. A decrease in *LDHA* (log2FC = -0.996), encoding lactate dehydrogenase A, in KG-1 cells further confirms suppression of anaerobic respiration in these cells. In MV4;11 cells, we also found a decrease in the expression of *SLC2A3* (log2FC = -1.395), encoding the glucose transporter GLUT3. At the same time, we found an increase in the expression of the *SLC2A5* gene (log2FC = 0.845), encoding GLUT5, which is responsible for the transport of fructose.

To check what happens with the metabolic indices in AML cells after exposure to CDK8/19i, we also grouped the DEGs of all the studied AML and obtained the average values per group (DEGs indices) (Fig. 2E). Indeed, only in sensitive AML cell lines after 72h of incubation in the presence of SenB there is a decrease in the metabolic indices of glycolysis and the anaerobic module. In sensitive AML cells, we also detected a number of DEGs associated with mitochondrial functions. In MV4;11, genes encoding the *MTO1* (log2FC = 1.658) and *NARS2* (log2FC = 1.001) proteins, which are involved in mitochondrial gene translation and amino acid metabolism, were upregulated. At the same time, calcium transport genes (*MICU1*, *MCU*) were downregulated by 2-fold. In KG-1, we found a decrease in the ribosomal gene *MRLPL27* (log2FC = -0.995) and phosphoenolpyruvate carboxykinase 2 *PCK2* (log2FC = - 1.008). These data support our hypothesis that the sensitivity of AML cells to CDK8/19i is determined by their metabolic patterns. Notably no metabolic genes were affected cell lines resistant to CDK8/19i (Fig. 2D). Interestingly, transcriptional changes, as well as sensitivity, in AML cells are also independent of the level of CDK8 or CDK19 in the cells (Fig. 2C).

To identify potential CDK8/19 targets contributing to the CDK8/19i-sensitivity among transcription factors (TF), we analyzed the DEGs of all cell lines using the decoupleR library in R (Fig. 2F-H, Supplementary Fig. S6). We found that 72 h after SNX631 treatment, KG-1 cells displayed a marked suppression of HIF1A transcriptional activity (ULM score = -5.58). In contrast, THP-1 cells showed an increase in HIF1A activity (ULM score = 3.58), suggesting that THP-1 cells retain the capacity to mount a hypoxic or pseudo-hypoxic adaptive response, unlike KG-1. No significant HIF1A enrichment was observed in either the most sensitive MV4;11 or the most resistant Kasumi-1 cells.

In both sensitive cell lines, KG-1 and MV4;11, we observed a more than 4-fold reduction in the transcriptional activity of SP1/SP3 target genes and nearly 3-fold reduction of SMARCA1/4 target genes, which was absent in the resistant THP-1 and Kasumi-1 lines. SP1 and SP3 share overlapping functions and regulate the activity of GC-box-binding elements critical for a broad repertoire of metabolic genes [21–24]. Although we did not identify substantial overlap in DEGs between the resistant cell lines, THP-1 and Kasumi-1 shared enrichment of 20 common TFs (Fig. 2G), the majority of which are associated with RNA polymerase II transcriptional regulation and super-enhancer activity (Fig. 2H). It is possible that this may reflect epigenetic and transcriptional reprogramming in resistant cells THP-1 and Kasumi-1 that does not occur in sensitive KG-1 and MV4;11 cells.

### The sensitivity of AML cells to CDK8/19i is determined by their metabolic activity

To assess baseline metabolic activity in AML cells and its association with CDK8/19i response, we calculated metabolic gene set indices in untreated samples (basal transcription), using the same approach as for DepMap expression data similar to those for the DepMap expression data. We identified gene groups that correlate with the sensitivity of AML cells to CDK8/19i, and they were nearly identical to the results of the DepMap expression data analysis: the Krebs cycle (TCA), the anaerobic respiration gene group (Anaerobic_module), and ETC complexes I and II (OXPHOS_I, OXPHOS_II) (Fig. 3A, upper). In addition, we found that the values of the basal transcription indices of OXPHOS (except for complex III), the anaerobic respiration and the TCA correlate with the sensitivity of cells to CDK8/19i as determined by resazurin staining.

**Figure 3.**
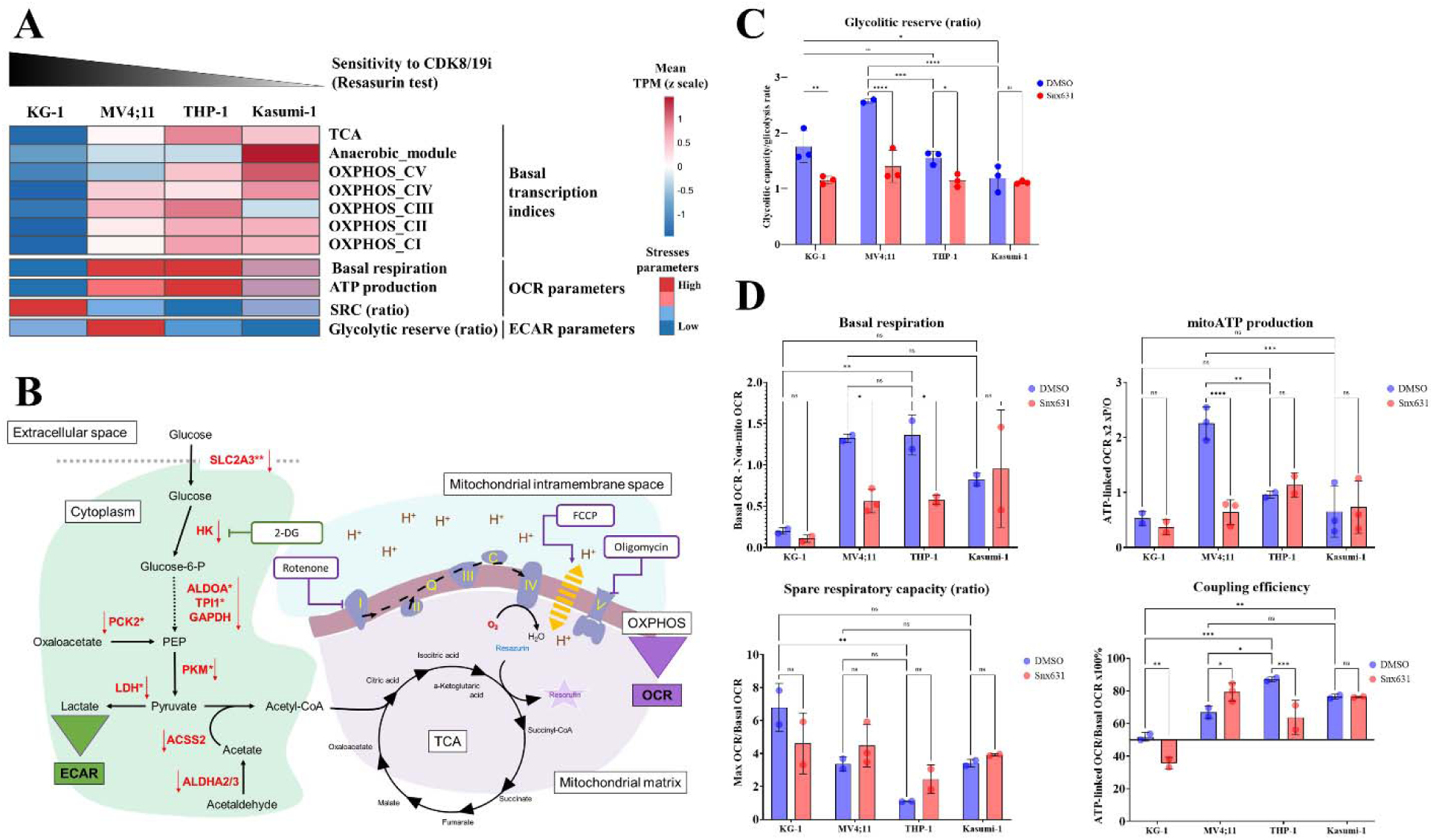
The sensitivity of AML cells to CDK8/19i directly correlates with metabolic indices and mitostress parameters. **A.** Heatmap of metabolic indices and main MitoStress and GlycoStress parameters obtained using the metabolic assay (OCR and ECAR respectively). **B.** A general diagram of the relationship between glycolysis (green area), the Krebs cycle (TCA) and OXPHOS (purple area). Genes whose expression is reduced in KG-1*, MV4;11**, or both lines after 72h in the presence of SenB, as determined by RNA sequencing, are highlighted in red. Inhibitors added during OCR or ECAR assays are shown in white squares with green or purple frames respectively. The diagram shows that resazurin is primarily reduced in the mitochondria during succinyl-CoA synthesis. **C.** The influence of 1 µM SNX631 on the glycolytic capacity of AML cells. The following metabolic parameters were calculated as follows: glycolysis = highest ECAR after glucose injection – lower ECAR after 2-DG injection, glycolytic capacity = highest ECAR after oligomycin injection − lower ECAR after 2-DG injection, Glycolytic reserve (ratio) = glycolytic capacity/glycolysis. Data are presented as mean ± SD. Statistical analysis was performed using Two-way ANOVA with Bonferroni correction. *ns* – non-significant, * – *p*<0.5, ** – *p*<0.1, *** – *p*<0.01. **D.** The influence of 1 µM SNX631 on the OXPHOS parameters of AML cells. The following metabolic parameters were calculated as follows: non-mitochondrial respiration (NMR) = lowest OCR after the injection of rotenone and antimycin A, basal respiration = max OCR before the oligomycin injections - NMR, proton leak = lowest OCR after the oligomycin injection, ATP-linked respiration = basal respiration - proton leak, maximum respiratory capacity (MRC) = highest OCR after the FCCP injection - NMR, spare respiratory capacity (SRC) ratio = MRC/basal respiration, mitoATP production = ATP-linked respiration x2 xP/O (standard ratio by Agilent is 2.5), coupling efficiency = mitoATP production/basal respiration. Data are presented as mean ± SD. Statistical analysis was performed using Two-way ANOVA with Bonferroni correction. *ns* – non-significant, * – *p*<0.5, ** – *p*<0.1, *** – *p*<0.01, **** – *p*<0.001.

To test whether the sensitivity of AML to CDK8/19i depends on their basal energy metabolism rate, we assessed their mitochondrial (oxygen consumption rate; OCR) and glycolytic respiration (extracellular acidification rate; ECAR) in the presence and absence of more active CDK8/19i SNX631 (Fig. 3, Supplementary Fig. S7). Measurement of GlycoStress parameters (the rate of glycolytic acidification of the medium after sequential addition of oligomycin and 2DG in the presence of high glucose concentrations) (Fig. 3B) in AML cells after addition of 1 μM SNX631 showed that MV4;11 cells have the most pronounced glycolytic reserve, which represents the capacity of a cell to increase its rate of glycolysis in response to sudden energy demands. Decrease in glycolytic reserve in response to CDK8/19i is observed only in sensitive KG-1 and MV4;11 cells (Fig. 3A-C), which is consistent with RNA-sequencing data (Fig. 3B, red).

MitoStress parameters (measurement of the rate of oxygen consumption in the presence of oligomycin, FССP and a combination of rotenone and actinomycin) were calculated based on the rate of cellular OCR (Fig. 3B,D, Supplementary Fig. S8A,B), which normally occurs during OXPHOS. We found that the most sensitive cells, as determined by resazurin staining (KG-1), had lower values for all calculated MitoStress parameters, except for the spare respiratory capacity (SRC), which characterizes unused mitochondrial potential (Fig. 3D, Supplementary Fig. S8B). Taking into account the low metabolic index values, we can conclude that KG-1 utilize mitochondria poorly and inefficiently (the lowest coupling efficiency values (which indicates efficiency of oxygen consumption for ATP synthesis) among all four lines ∼50%). MV4;11, which demonstrated the highest cell death when exposed to CDK8/19i, exhibited higher mitochondrial ATP production (mitoATP production) and basal mitochondrial respiration (oxygen consumption in basal conditions). MV4;11 also exhibited the highest proton leak values (the passive movement of protons across the inner mitochondrial membrane back into the matrix) and higher than KG-1 coupling efficiency values (under 70%). Thus, although MV4;11 utilized mitochondria to a greater extent than KG-1, they utilized it less efficiently than CDK8/19i-resistant THP-1 (coupling efficiency 90%), which had similar basal respiration values. The most resistance to CDK8/19i Kasumi-1 cells have moderate basal respiration and SRC values, low proton leak, and sufficient coupling efficiency (up to 80%). The calculated parameters are also consistent with the indices and cell sensitivity to CDK8/19i (Fig. 3A). Our data indicates that the sensitivity of AML cells is most correlated with their glycolytic reserve (Fig. 3A, lower panel).

After 72 h of treatment with SNX631, mitoATP production, basal respiration and proton leak decreased more than 2-fold in MV4;11 cells, while coupling efficiency increased up to 80% (Fig. 3D, Supplementary Fig. S8B). These data indicate a slowdown of mitochondrial function in the MV4;11 after SNX631 treatment. However, in KG-1 cells, we found only a decrease of coupling efficiency values by almost 25% (Fig. 3C), although there were no significant changes in the main parameters of MitoStress. In CDK8/19i-resistant THP-1 cells, we detected major changes only in basal respiration (2-fold decrease), which also resulted in a decrease in the coupling efficiency parameter. In Kasumi-1 cells after 72 h in the presence of SNX631 we found no changes in mitochondrial respiration parameters. Thus, according to the data obtained, the most significant changes in mitochondrial respiration ang glycolysis occur in the most sensitive to CDK8/19i AML cells.

### CDK8/19 inhibition leads to a decrease in glutamine metabolites in CDK8/19i sensitive AML cells

To confirm that CDK8/19i leads to metabolic changes in AML cells, we performed a GC-MS metabolomic analysis on all studied cell lines 72 h after SNX631 treatment (Fig. 4A, Supplementary Fig. S2). Consistent with our hypothesis, significant reduction in the metabolite levels was observed only in the CDK8/19i-sensitive KG-1 and MV4;11 cell lines. In the resistant THP-1 and Kasumi-1 lines the number of decreased metabolites was absent.

**Figure 4.**
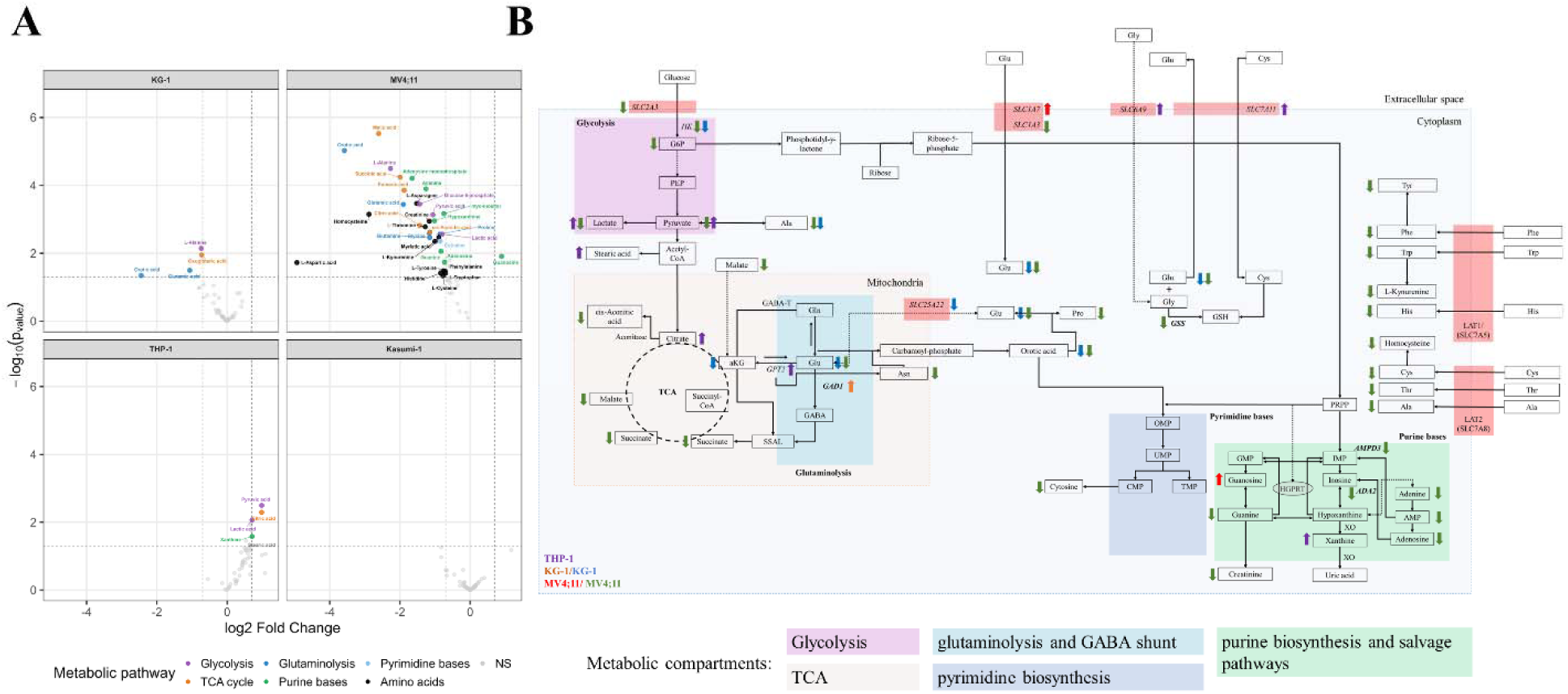
CDK8/19 inhibition leads to metabolic disorders only in sensitivity AML cell lines. **A.** Distribution graphs (Volcano plots) of differentially metabolic concentration after 72h of incubation of AML cells in the presence of 1 μM SNX631. **B.** Pathway map illustrating the metabolic changes observed in KG-1, MV4;11, and THP-1 cells 72 h after treatment with the CDK8/19 inhibitor SNX631. Arrows indicate the direction of change in metabolite levels or gene expression (↓ decreased, ↑ increased) for the indicated cell line, color-coded as follows: purple for THP-1, orange or blue - KG-1, red or green - MV4;11. DEGs are shown in italics; highlighted gene labels (light red background) indicate transporters with changed expression. Dashed arrows denote multi-step reactions. Abbreviations: αKG, α-ketoglutarate; G6P, glucose-6-phosphate; PEP, phosphoenolpyruvate; GABA, γ-aminobutyric acid; Ala, alanine; Asn, asparagine; Cys, cysteine; Gln, glutamine; Glu, glutamate; Gly, glycine; His, histidine; Phe, phenylalanine; Pro, proline; Thr, threonine; Trp, tryptophan; Tyr, tyrosine; GSH, glutathione; OMP, orotidine-5′-monophosphate; UMP, uridine monophosphate; IMP, inosine monophosphate; AMP, adenosine monophosphate; GMP, guanosine monophosphate; ADA2, adenosine deaminase 2; AMPD3, adenosine monophosphate deaminase 3; GABA-T, GABA transaminase; GPT2, glutamic-pyruvic transaminase 2; HGPRT, hypoxanthine-guanine phosphoribosyltransferase; XO, xanthine oxidase.

According to GC-MS data, we found that in both sensitive AML cell lines MV4;11 and KG-1 there are a decrease in TCA and glutaminolysis metabolites (Fig. 4A). Metabolomic data analysis revealed a nearly 2-fold decrease in major glutamine derivatives (glutamic acid and α-ketoglutarate (αKG)) in KG-1 cells, alongside an almost 5-fold decrease in orotic acid, a common nucleotide precursor (Fig. 4B). MV4;11 cells, which were the most sensitive based on cell cycle analysis, exhibited even more profound metabolic disruptions after 72 h. In addition to a more than 2-fold decrease in glutamine and glutamic acid, and a 12-fold decrease in orotic acid, MV4;11 cells showed significantly suppressed TCA metabolites and depleted adenine, cytosine, and guanine. Notably, the synthesis of all these depleted metabolites directly or indirectly depends on αKG as a precursor and glutamine as a nitrogen donor. Furthermore, several essential amino acids and their derivatives were significantly reduced in MV4;11 cells, suggesting the activation of alternative catabolic pathways. As previously described, SenB treatment for 72 h downregulates *HK3* and *SLC2A3* expression in MV4;11 cells (Fig. 2D). This downregulation may explain the significant decrease in glucose-6-phosphate (G6P) and pyruvate observed in our metabolomic analysis (Fig. 4B). In THP-1 cells, which possess moderate resistance to CDK8/19i, we found a nearly 2-fold increase in pyruvate, lactate, citrate, and stearic acid. As depicted in Fig. 4B, this metabolite accumulation is consistent with TCA cycle decoupling[25]. Additionally, THP-1 cells exhibited a 1.6-fold increase in xanthine levels, likely driven by the constitutively high xanthine oxidase expression previously described in this cell line[26]. The most resistant Kasumi-1 cells showed no significant changes in their metabolic profile. Overall, these data confirm that the extent of metabolic disruption directly correlates with cell line sensitivity to CDK8/19i.

## Discussion

We hypothesized that CDK8/19-sensitive AML cell lines differ from resistant cells primarily by their metabolic profile and the transcriptomic changes caused by CDK8/19i associated with both glycolysis and mitochondrial function. Notably, neither sensitivity nor the transcriptional response to CDK8/19i correlated with CDK8 or CDK19 protein levels (Fig. 2C). Glycolytic genes were notably downregulated only in sensitive AML lines, with resistant cells exhibiting no changes (Fig. 2D,E). In MV4;11, decreased *GLUT3* and *HK* expression was accompanied by reduced G6P levels, consistent with HCT116 cells exposed to CDK8/19i[15]. As was shown, HIF1A orchestrated transcription appeared to be dependent from CDK8, especially in hypoxic condition through promoter-proximal pause release mechanism. SNX631 treatment caused a decrease in key MitoStress parameters predominantly in sensitive cells, while in THP-1 only basal respiration was reduced, and Kasumi-1 parameters remained unchanged relative to controls after 72 h (Fig. 3D). Coupling efficiency below 80% (observed in both KG-1 and MV4;11) indicates inefficient mitochondrial energy transduction only in the sensitive lines (Supplementary Fig. S8B). Based on SRC ratio data, Kasumi-1 displays greater dependence on mitochondrial than on glycolytic activity, consistent with its highest metabolic index values across all lines (Fig. 3A,D, Supplementary Fig. S8) and previous findings[27]. Although MV4;11 and THP-1 behave similarly in the OCR assay, MV4;11 has lower basal metabolic index values (Fig. 3A, upper). Previous studies indicating that THP-1 is more OXPHOS dependent than KG-1 and MV4;11 [28–32], collectively supporting our hypothesis that a predominantly glycolytic baseline metabolic state confers greater sensitivity to CDK8/19 inhibition.

Transcriptome analyses identified gene sets correlating with AML sensitivity to CDK8/19i: notably, genes of the Krebs cycle, anaerobic respiration, and ETC complexes I and II consistent with analysis of expression DepMap data (Fig. 1D,E). Metabolic indices calculated from DEG profiles decrease only in sensitive cell lines during inhibitor exposure (Fig. 2E). Furthermore, sensitivity of AML cells to CDK8/19i correlated most strongly with dependency on the KO of metabolism-associated genes (Fig. 1C). Our data suggest that AML sensitivity to CDK8/19i is determined not by a single molecular pathway, but rather by the metabolic landscape.

GO enrichment data (Supplementary Fig. S4B) are consistent with the study by Pelish et al., in which CA inhibition after 6 h also resulted in changes in the expression of gene groups related to leukocyte activation and the inflammatory response[4]. The authors show that in AML cells, in response to CA, there is an overexpression of genes encoding super-enhancer associated transcription factors, such as *CEBPA*, *IRF8*, *IRF1*, and *ETV6*. We analyzed our DEGs for the presence of these genes in all studied AML lines and found an increase in the expression of the *ETV6*, *IRF1*, and *IRF8* genes in MV4;11 cells and only *IRF1* in Kasumi-1 cells (Supplementary Table 2). Moreover, the expression of the *IRF1/8* genes was increased in both cell lines by 2 or more times. However, these changes are found in resistant cells too and they do not explain the cell death mechanism.

Evidence is emerging that CDK8/19 modulate cellular metabolism, including mitochondrial function and glycolysis[14,15,33–36]. CDK8, to a greater extent than CDK19, is involved in the mechanisms regulating the cellular response to starvation, both through the transcriptional regulation of early hypoxia response genes, as a cofactor of HIF1A[14,15,37], and through the phosphorylation of metabolism-associated transcription factors, such as SREPB1c[37]. Moreover, CDK8 is negatively regulated by the important metabolic regulator mTOR[13,37]. Interestingly, both loss and overexpression of CDK8 can lead to a decrease in the expression of glycolytic and mitochondrial genes[34]. The differential engagement of downstream signaling pathways in sensitive and resistant AML cell lines (JAK/STAT in MV4;11 or PI3K/Akt and MAPK in Kasumi-1), likely reflecting their distinct molecular backgrounds may further contribute to the different metabolic responses observed upon CDK8/19i[38]. Notably, these differences in response could not be attributed to CDK8 or CDK19 protein levels, which were comparable across all cell lines (Fig. 2C).

In both sensitive cell lines, KG-1 and MV4;11, enrichment of TFs be RNA-seq data shown a nearly 3-fold reduction in activity of SMARCA1/4 – core components of the SWI/SNF chromatin remodeling complex (Fig. 2F). This is consistent with the previously demonstrated physical interaction between the CDK8/19-Mediator kinase module and SWI/SNF components and regulates enhancer-mediated transcriptional output of specific targets[17,39–41]. CDK8/19i-resistant AML cells, THP-1 and Kasumi-1, shared enrichment of 20 common TFs are associated with RNA polymerase II transcriptional regulation and super-enhancer activity (Fig. 2G,H). These data indicates that epigenetic reprogramming occurs in all AML cells after CDK8/19i treatment.

In both CDK8/19i-sensitive cell lines we observed deficit of glutamine and its derivatives after 72 h SNX631 treatment (Fig. 4). We hypothesize that suppression of HIF1A-dependent transcription, including genes encoding glycolytic enzymes, upon CDK8/19 inhibition, via the mechanism described in[15], reduces glycolytic flux and forces cells to rely more heavily on glutamine as an alternative carbon and nitrogen source. This increased glutamine consumption, combined with likely disruption of the TCA cycle, leads to progressive depletion of αKG, since glutamine-derived glutamate represents the primary anaplerotic route for αKG replenishment when TCA activity is impaired. Notably, KG-1 cells appear capable of compensating for reduced αKG through 2-fold upregulation of *GAD1*, which encodes glutamate decarboxylase and enables the GABA shunt, an alternative pathway that converts αKG to succinate via GABA, bypassing canonical TCA reactions[42]. This interpretation is supported by unchanged succinate levels despite decreased αKG in KG-1 cells, suggesting that the TCA cycle remains functionally maintained through this compensatory route. Transcriptional activity of HIF1A target genes is suppressed in KG-1 cells following CDK8/19i (Fig. 2F), in accordance with that reduced αKG levels would be expected to stabilize HIF1A protein under normoxic conditions[43]. In MV4;11, the cell line most sensitive to CDK8/19i, TCA disruption and the ensuing compensatory glutamine consumption likely lead to a broader metabolic collapse. As one of the principal nitrogen and carbon donors for biosynthetic reactions, glutamine depletion in MV4;11 leads to concurrent decreases in precursors for purine and pyrimidine biosynthesis, including orotic acid, whose synthesis depends on glutamine-derived carbamoyl phosphate, as well as multiple essential amino acids, indicating an inability to activate any effective compensatory mechanism in these cells. The resistant cell lines displayed a completely different metabolic response. In Kasumi-1, the most resistant line, no significant metabolic changes were detected following SNX631 treatment. In THP-1 cells, which exhibit moderate resistance, we observed increased levels of pyruvate, lactate, citrate, and stearic acid, indicating a metabolic state in which glycolytic flux is preserved or enhanced while mitochondrial oxidation is partially redirected toward lipogenesis (Fig. 4). Excess pyruvate supports lactate production via lactate dehydrogenase and fuels de novo fatty acid synthesis via acetyl-CoA, with citrate accumulation reflecting its overflow into the proximal TCA cycle. The accumulation of lower glycolytic metabolites (Fig.1E, Fig4B) appears to be related to TCA cycle uncoupling and slower metabolism, as shown in Fig. 2A. Thus, in all AML cells except Kasumi-1, CDK8/19i lead to TCA cycle disruptions, but only sensitive AML cells are unable to fully compensate for these disruptions. HIF1A has been shown to regulate the shift of glutamine metabolism from oxidative stress to reductive carboxylation, which supports lipid synthesis and tumor growth under oxygen deficiency[44]. For example, HIF1A reduces cellular damage by positively regulating *SLC7A11*, which increases antioxidant capacity in neurons[45]. The 4-fold increase of *SLC7A11* in THP-1 and activation of HIF1A-dependent transcription can be caused by the compensatory mechanism of pseudo-hypoxia in response to low levels αKG[43]. Furthermore, compensatory transcriptional upregulation of *GPT2*, *SLC6A9*, and *SLC7A11* suggests that THP-1 cells actively maintain glutamate homeostasis under inhibitor treatment, providing an additional mechanism of metabolic resilience. The absence of disturbances in glutaminolysis in resistant cells indicates that the decrease in glutamine in sensitive cells is an exclusively compensatory mechanism. These data support the results of previous studies indicating that Kasumi-1 cells are most sensitive to inhibition of glutaminolysis [46].

Collectively, these findings demonstrate that CDK8/19 play a key role in sustaining glutaminolysis and broader metabolic homeostasis in AML cells, and that sensitivity to CDK8/19i is determined by the intrinsically glycolytic baseline metabolic profile of the cell.

## Acknowledgments

We thank Dr. V. V. Grinev for Kasumi-1 cells. This work was performed using the equipment of IGB RAS facilities supported by the Ministry of Science and Higher Education of the Russian Federation.

## Contributions

VEA., BAV and TVV analyzed the data and wrote the manuscript. VEA, ADV, DAA, MEV and PNG carried out experiments. BAV and TVV acquired funding, supervised the research, and wrote the manuscript.

## Abbreviations

AML: acute myeloid leukemia
CDK8/19: cyclin-dependent kinases 8/19
CDK8/19i: CDK8/19 inhibitors
ECAR: extracellular acidification rate
ETC: electron transport chain
KO: knockout
minIC_50_: minimal published IC_50_
OCR: oxygen consumption rate
OXPHOS: oxidative phosphorylation
SenB: senexin B
SRC: spare respiratory capacity
TF: transcription factor
αKG: α-ketoglutaric acid

