## Supplementary figures and images for "The sensitivity of acute myeloid leukemia to CDK8/19 inhibitors is determined by their metabolic profile"

### Supplementary figure S1

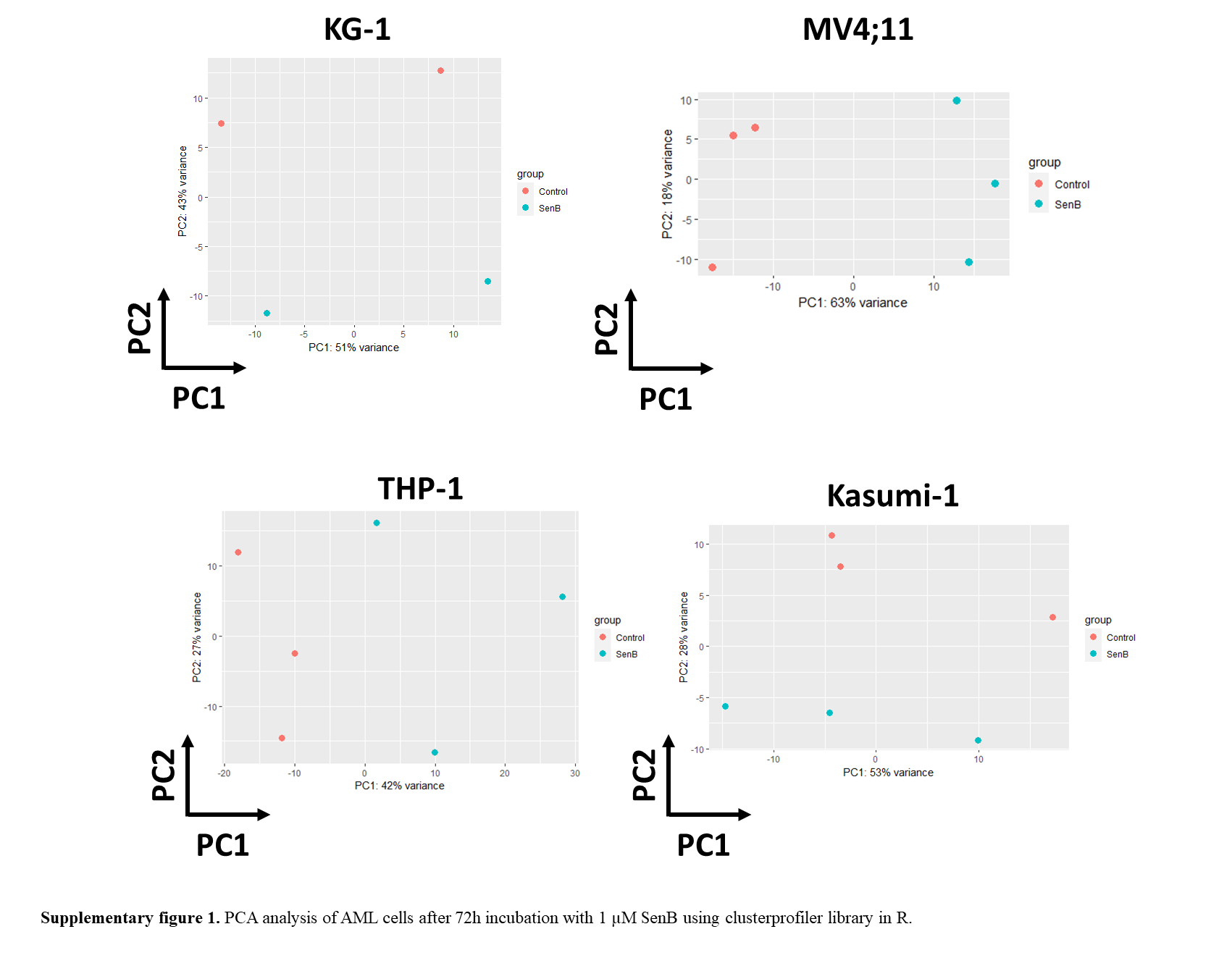

### Supplementary figure S2

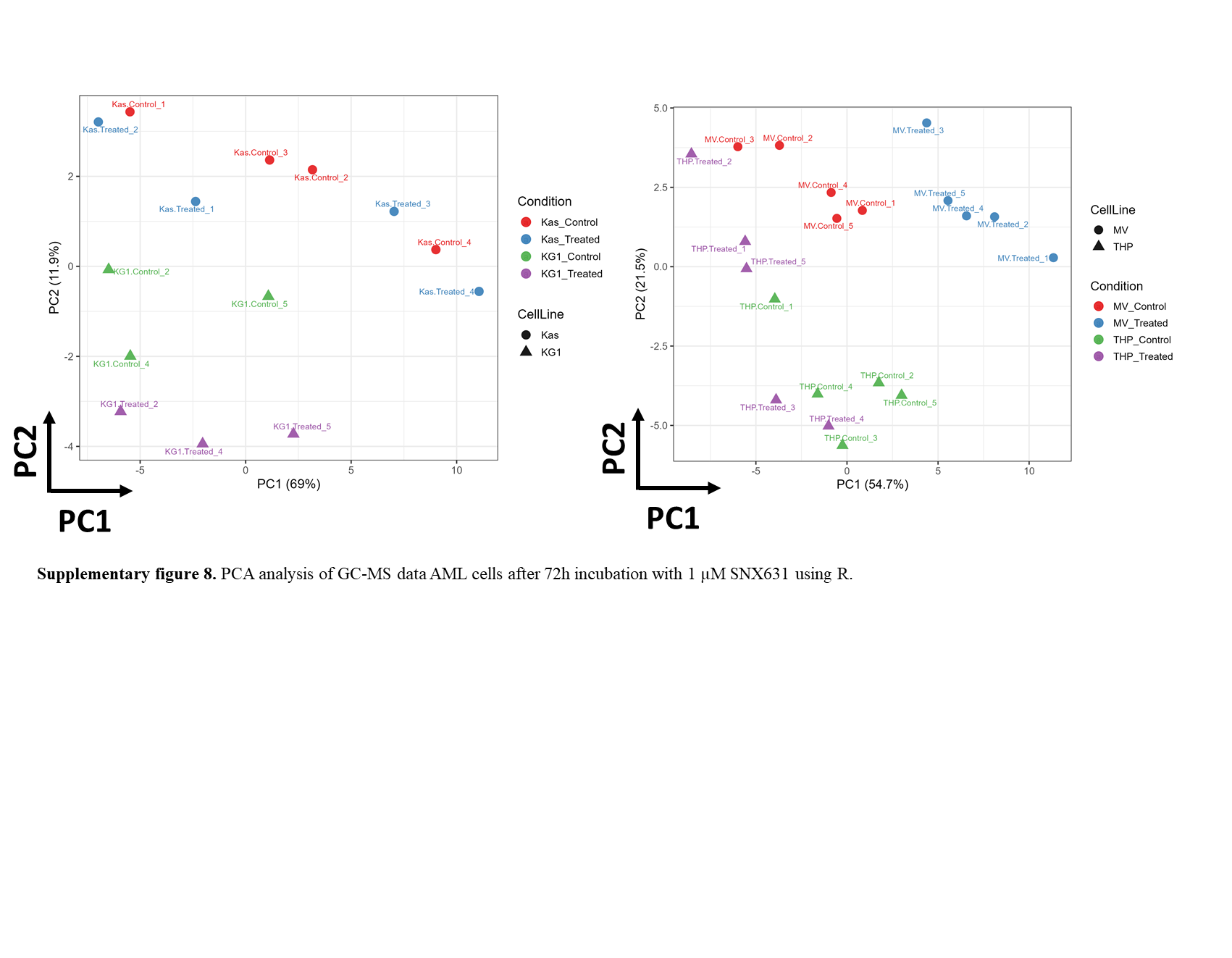

### Supplementary figure S3

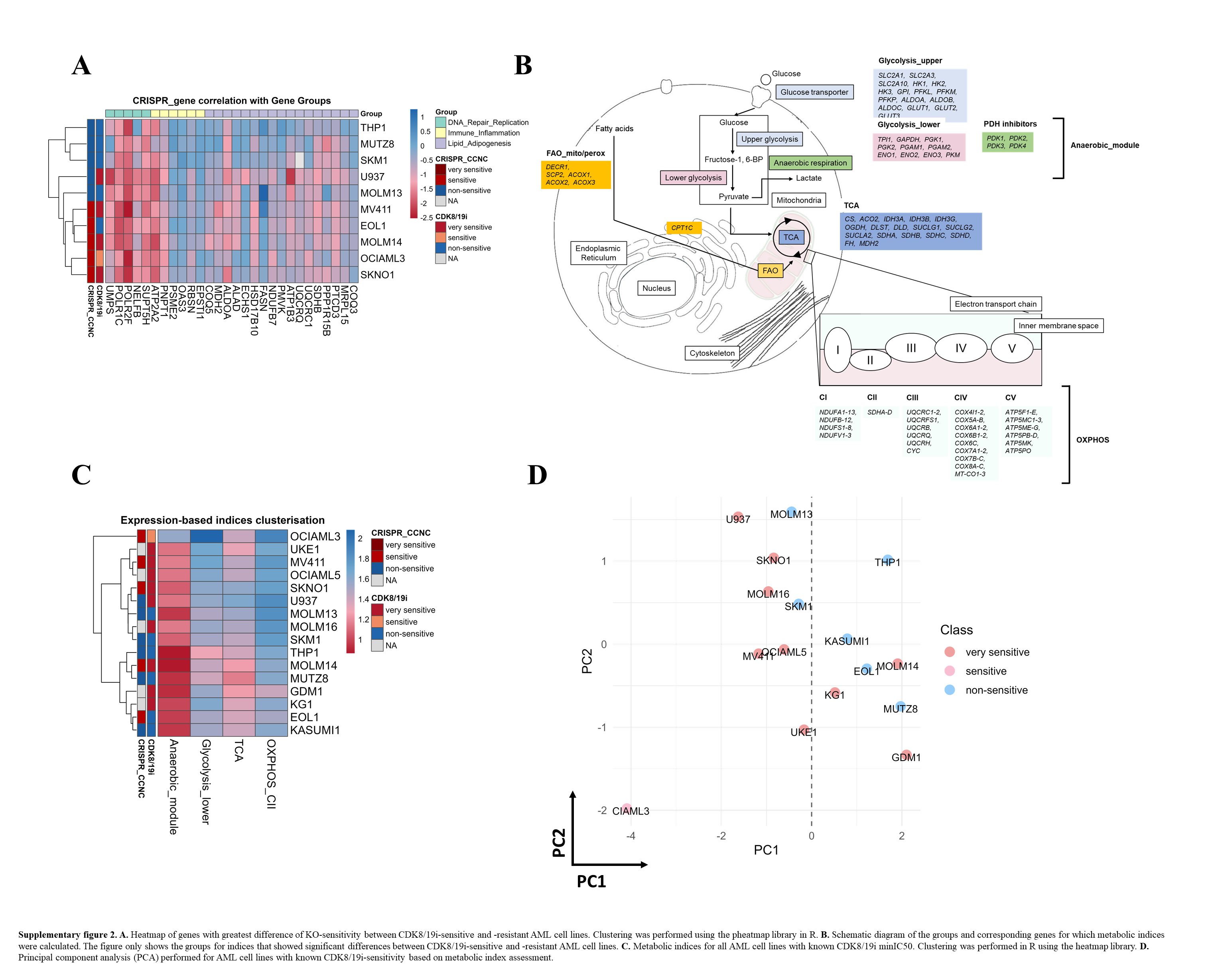

### Supplementary figure S4

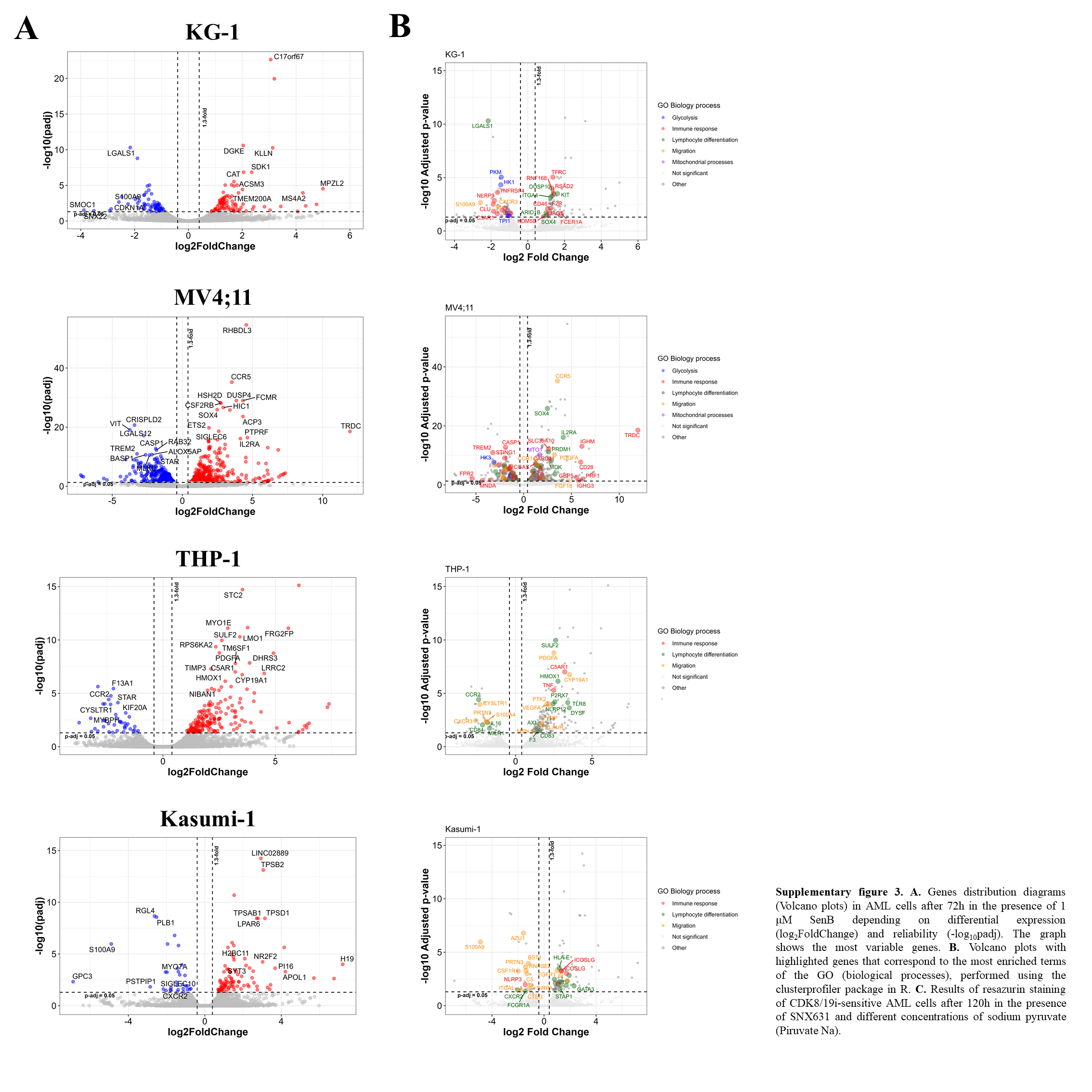

### Supplementary figure S5

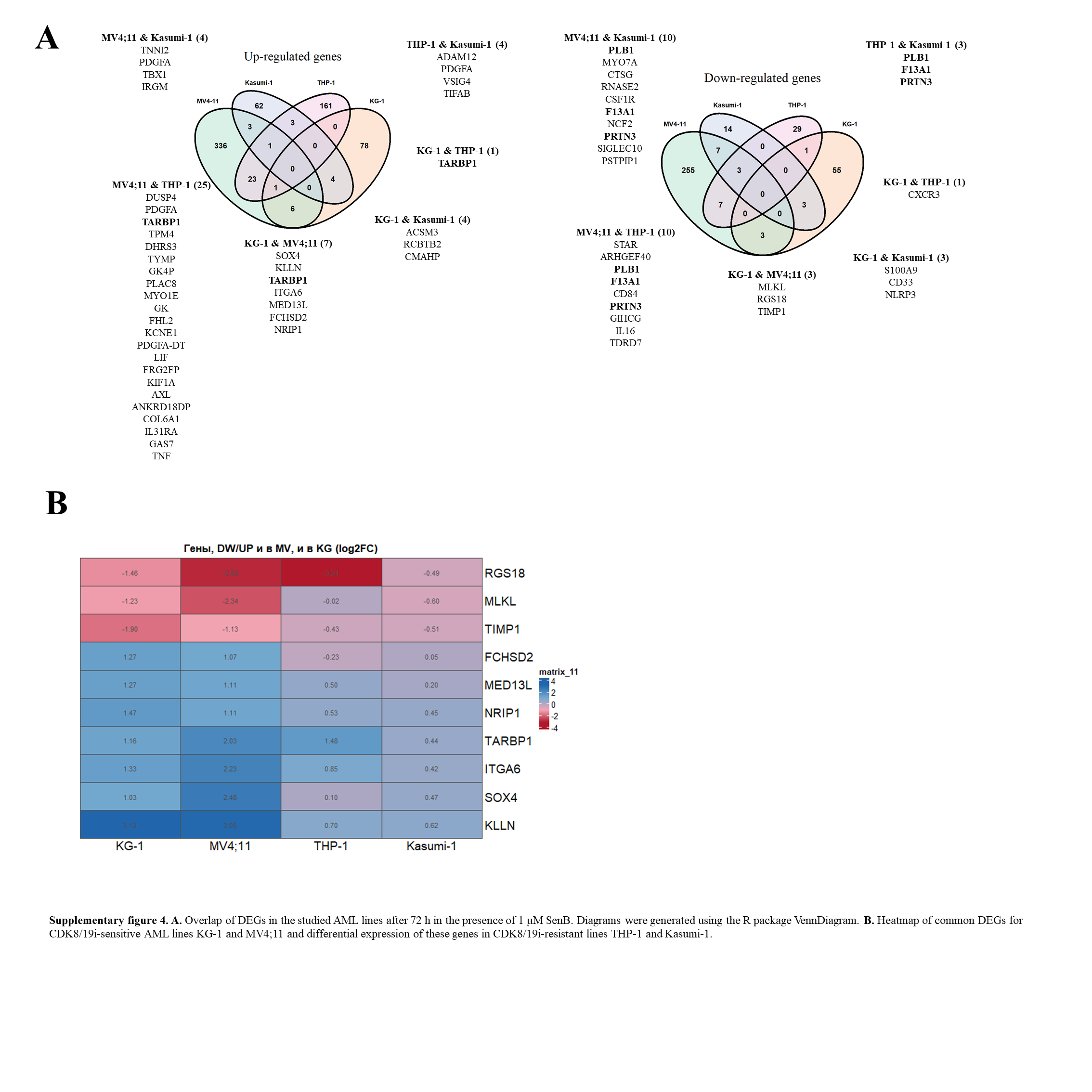

### Supplementary figure S6

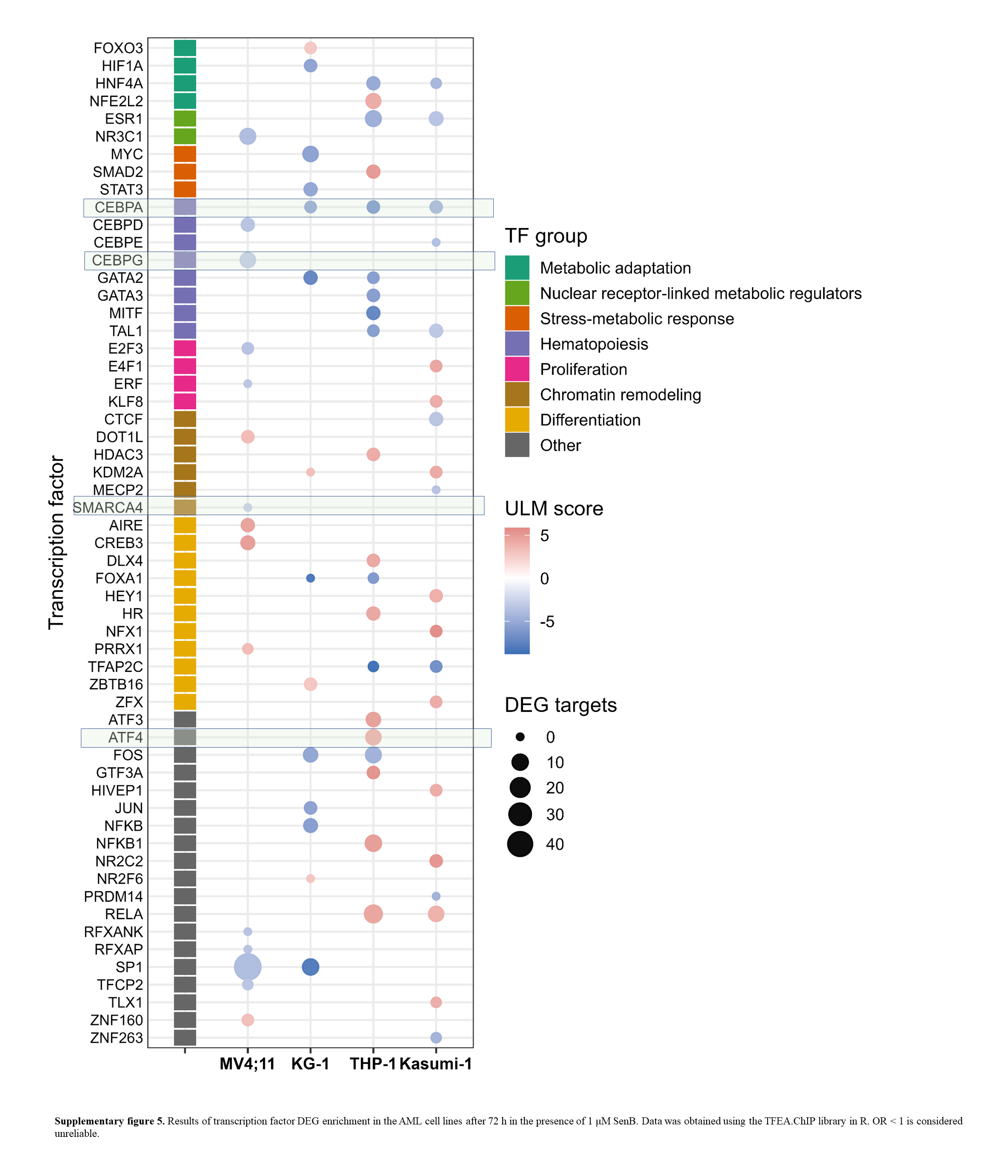

### Supplementary figure S7

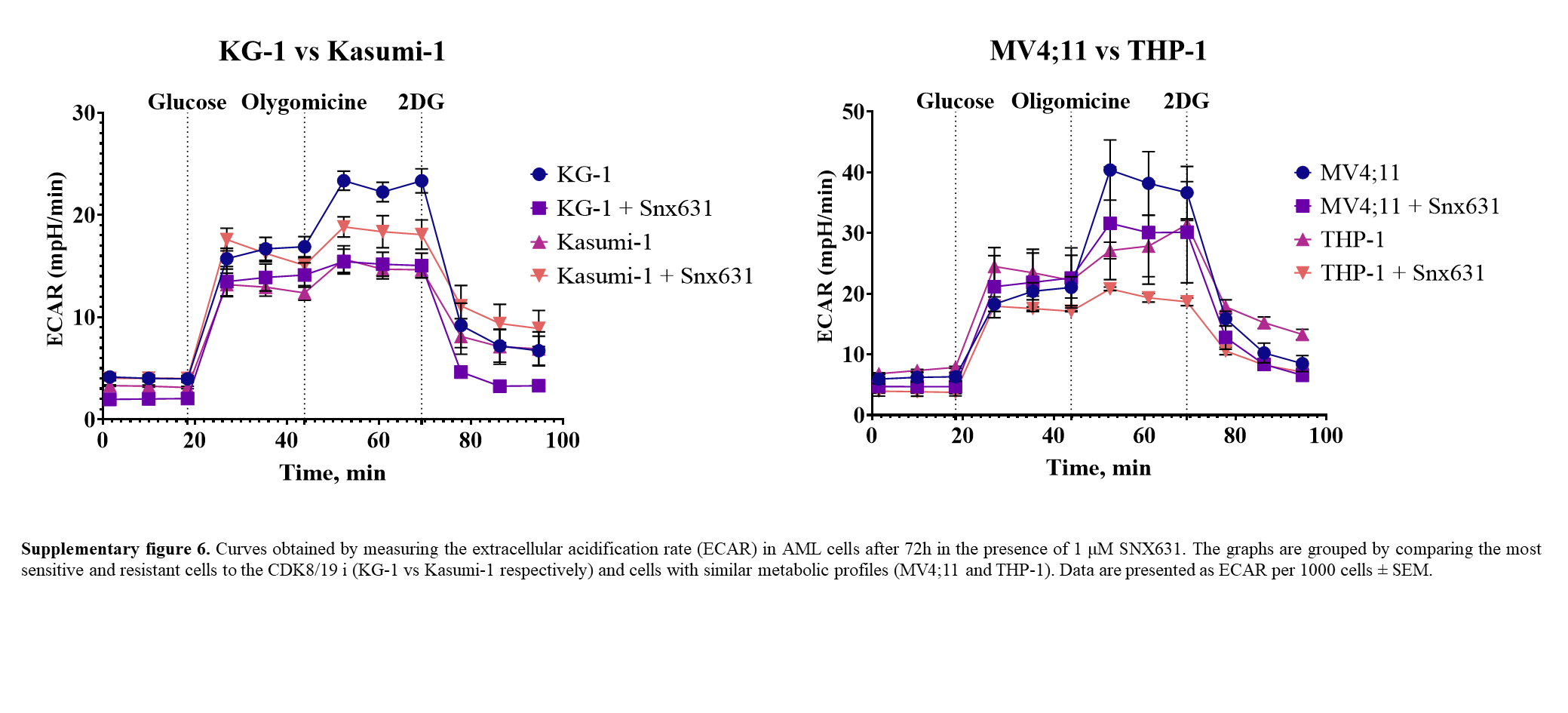

### Supplementary figure S8

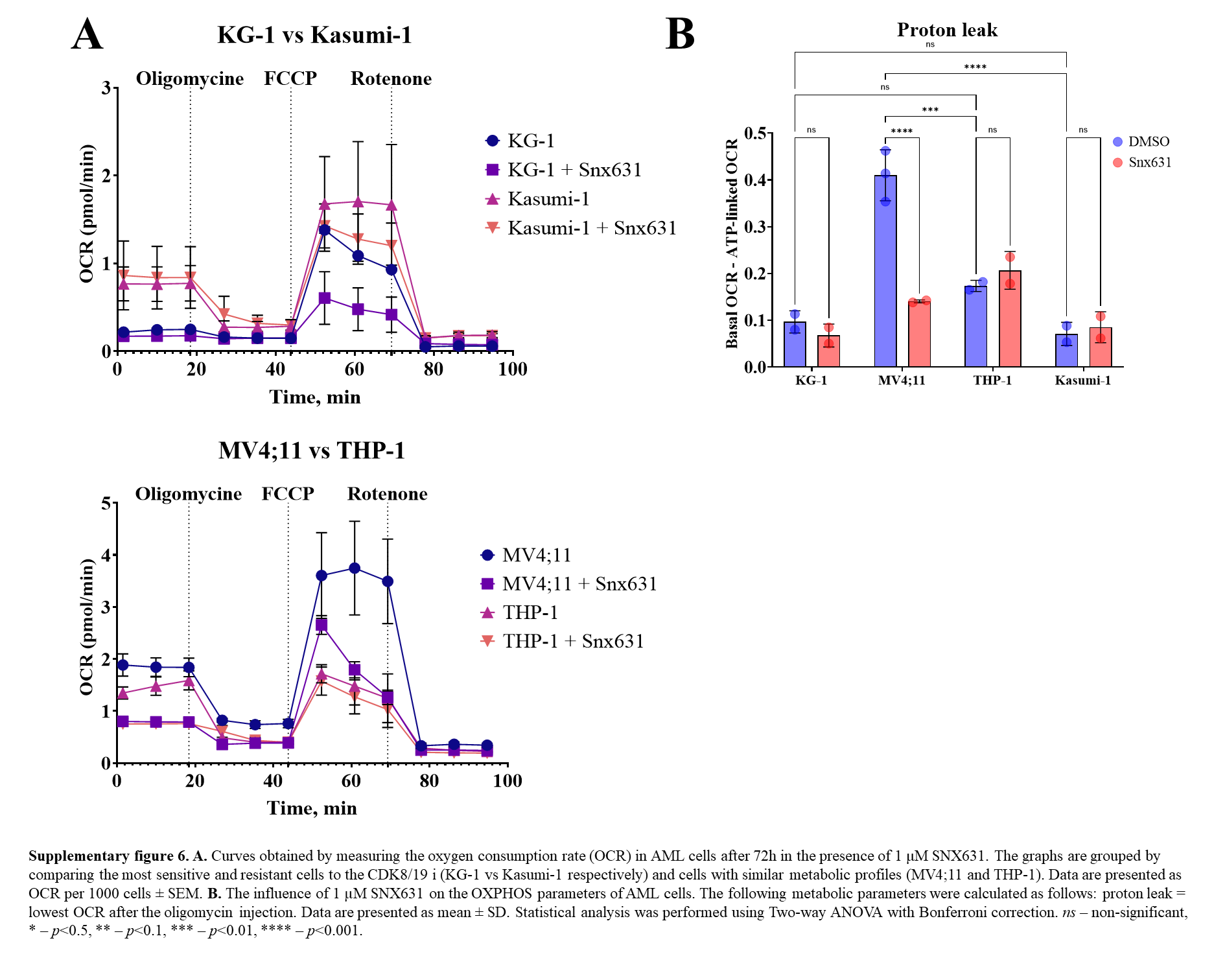
